# Chemotherapeutic Nanoparticles Accumulate in the Female Reproductive System during Ovulation Affecting Fertility and Anticancer Activity

**DOI:** 10.1101/2020.07.22.216168

**Authors:** Maria Poley, Yael Shammai, Maya Kaduri, Lilach Koren, Omer Adir, Jeny Shklover, Janna Shainsky, Irit Ben-Aharon, Assaf Zinger, Avi Schroeder

**Author notes:** To whom correspondence should be addressed: A.Z., A.S.

## Abstract

Throughout the female menstrual cycle, physiological changes occur that affect the biodistribution of nanoparticles within the reproductive system. This can have positive or negative effects. We demonstrate a 2-fold increase in nanoparticle accumulation in the ovaries during female mouse ovulation compared to the non-ovulatory stage following intravenous administration. Accumulation in the reproductive system is favored by nanoparticles smaller than 100 nm. Chemotherapeutic nanoparticles administered during ovulation increased ovarian toxicity and decreased short-term and long-term fertility when compared to the free drug. Breast cancer treated with nanomedicines during ovulation results in higher drug accumulation in the reproductive system rather than at the site of the tumor, reducing treatment efficacy. Conversely, ovarian cancer treatment was improved by enhanced nanoparticle accumulation in the ovaries during ovulation. Our findings suggest that the menstrual cycle should be considered when designing and implementing nanotherapeutics for females.

## Introduction

The effect of the menstrual cycle on nanoparticle biodistribution and activity is overlooked in the study of therapeutic nanotechnologies. The female reproductive system, which harbors the egg reservoir, undergoes cyclic hormonal and physiological changes that lead to ovulation^1^. During the pre-ovulatory stage, increased blood flow, angiogenesis and perfusion within the reproductive system support oocyte maturation, with consequences for nanoparticle distribution (Figure 1a)^1,2^. For female mice, the estrous cycle (the equivalent of the human female menstrual cycle) is divided into four stages: diestrus, proestrus, estrus, and metestrus^3^. At the end of the proestrus stage, ovulation occurs, followed by the estrus stage.

**Figure 1.**
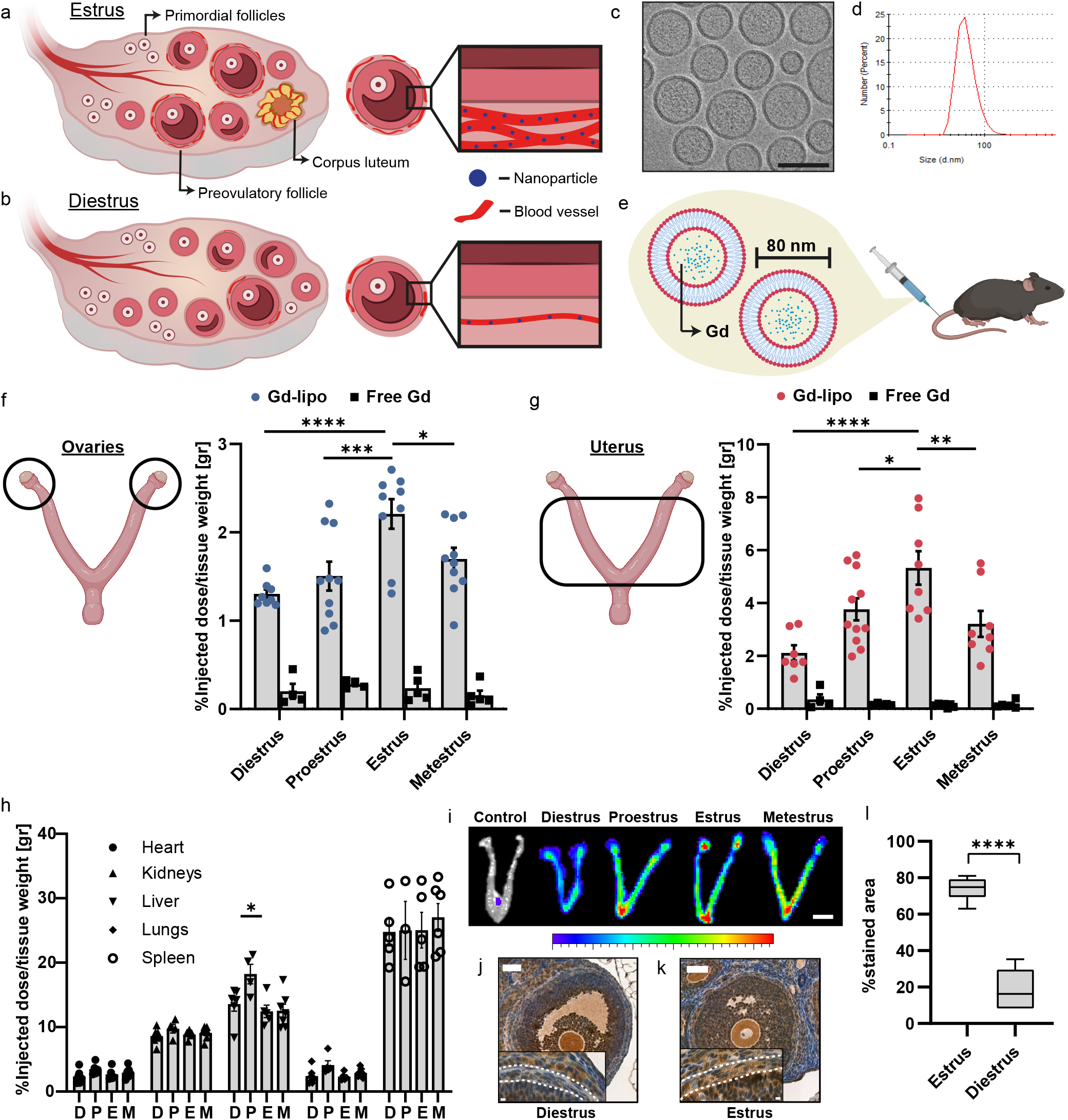
Biodistribution of nanoparticles during the female mouse menstrual cycle. During the estrus stage there is increased blood supply to the ovary to support preovulatory follicles. After ovulation, a dense blood network termed the corpus luteum is observed. Higher density of blood vessels around the follicle result in higher accumulation of nanoparticles (blue) in the reproductive system (a). By contrast, there are fewer blood vessels in the ovary and around the follicles specifically during the diestrus stage (b). We used 80-nm Gd liposomes characterized using Cryo-TEM images (c, scale bar - 100 nm) and DLS measurements (d). Gd-lipo or Free Gd were i.v. injected to female mice at different cycle stages (e) and their accumulation after 24 hours was quantified using elemental analysis. Results are shown as the percentage of Gd accumulated out of the total injected dose normalized to the organ’s weight. 2-fold more liposomes reached the ovaries at the estrus stage (n;9) compared to the diestrus stage (n;9) (f). 2.5-fold more liposomes reached the uterus at the estrus stage (n;8) compared to at the diestrus stage (n;7) (g). There was no differences in accumulation in other organs at different stages, excluding the liver (h). Ex-vivo fluorescent images of the female mouse reproductive system 24 hours post i.v. injection of 80 nm Cy5-labeled liposomes demonstrate the enhanced accumulation during the estrus stage (i, scale bar - 0.5 cm). Blood vessel density was evaluated using image analysis of immunohistochemistry staining of anti-CD31 at diestrus (j) and estrus stages (k) (scale bar - 100 μm) and was quantified as the percentage of stained area compared to a control stained only with secondary Ab (l). Results are shown as mean±SEM. Two-way ANOVA and Tukey’s t-test were used for statistical analysis of f-h, unpaired two-tail t-test was used for statistical analysis of l. *p<0.05, **p<0.01, ***p<0.001, ****p<0.0001.

Nanotechnologies are becoming important clinical tools, allowing accurate diagnosis and therapy^4–10^. Several nanotheraputic drugs are currently in clinical use for treatment of female-specific cancers, such as ovarian cancer^11,12^. Nanoparticle distribution to the female reproductive system can improve the efficacy of cancer treatments or pose a threat to fertility by causing ovarian toxicity and subsequent ovarian failure^13–16^.

There has been a recent increase in attention towards sex-specific medicine^17^. Foci include the study of female-specific administration routes, such as vaginal delivery^18–20^, and evaluation of sex differences in pharmacokinetic parameters of clinically-approved nanomedicines^21^. PEGylated liposomal doxorubicin has a slower overall clearance rate in females than in males^21^. The reason behind this sex-related difference is yet to be determined. It has recently been demonstrated that the cellular uptake of nanoparticles is dependent on the cell sex identity^22^. Preclinical characterization of nanoparticle biodistribution and pharmacokinetic parameters is therefore an important step in the development of nanocarriers, especially given the wide variety of nanoparticle types and applications^23–27^. Here, we study how the female menstrual cycle affects the accumulation of nanoparticles in the reproductive system. We then explore the effect of nanomedicines on female fertility and determine how the efficacy of nanomedicines is affected by the stages of the menstrual cycle.

## Results and Discussion

The female reproductive system undergoes cyclic physiological changes timed with the monthly menstrual cycle. One of these changes is the formation of new blood vessels around developing follicles before ovulation (Figure 1a, b). We studied the accumulation of nanoparticles in developing follicles before, during, and after ovulation using 80±10 nm liposomes loaded with gadolinium (Gd-lipo, Figure 1c, d) in female mice.

### Maximal nanoparticle accumulation in the reproductive system was measured during ovulation

The menstrual cycle of the female mouse is divided into four stages: diestrus, proestrus, estrus, and metestrus. The cycle stage was determined using vaginal cytology (Figure S1a). Gd-lipo or free Gd were injected intravenously (i.v.) (Figure 1e) to the tail vein of the mice at each of the four stages. 24 hours after the injection, accumulation of either Gd-lipo or free Gd in the ovaries (Figure 1f) and the uterus (Figure 1g) was quantified using elemental analysis. Maximal accumulation of Gd-lipo was assessed during the estrus stage for the ovaries (2.2%±0.17 of the injected dose, n=9) and the uterus (5.3% ± 0.6 of the injected dose, n=8). The lowest accumulation occurred during the diestrus stage, where only 1.25%±0.07 (n=9) and 2.1%±0.3 (n=7) of the injected dose reached the ovaries and the uterus, respectively. This amounts to a ~2-fold (*p*<0.0001) and ~2.5-fold (*p*<0.0001) increase in Gd-lipo accumulation during the estrus stage compared to the diestrus stage at the ovaries and uterus, respectively. The accumulation during the proestrus (1.5%±0.2, n=10 for ovaries, 3.8% ± 0.4 n=11 for uterus) and the metestrus (1.7%±0.1 n=10 for ovaries, 3.2%±0.5 n=8 for uterus) stages was significantly lower than during the estrus stage. In both the uterus and the ovaries, throughout the estrous cycle stages, the amount of free Gd compared to Gd-lipo was significantly lower (*p*<0.0001), indicating that the quantified Gd from the liposome-treated mice was due to liposomal accumulation. No significant differences in liposomal accumulation were found in the heart, lungs, kidneys, and spleen (Figure 1h) excluding the liver where a significant rise in liposomal accumulation was recorded during the proestrus stage compared to the diestrus, estrus or metestrus stages (*p*<0.05). This increase may be related to elevated estrogen levels during proestrus and its effect on cytochrome P450 liver metabolism^28^.

To further assess the enhanced nanoparticle accumulation during estrus, we visualized the reproductive tract 24 hours after an i.v. injection of 80 nm cy5-labeled liposomes (Figure 1i). The fluorescent intensity in the ovaries was highest during the estrus stage compared to the diestrus, proestrus, and metestrus stages (Figure 1i). These results are in agreement with the quantitative Gd-lipo biodistribution results shown above.

### Increased blood vessel density during the ovulatory stage

At the time of follicle development, as preparation for ovulation, new blood vessels are formed in the thecal layer surrounding the follicle^29^. To validate the increase in blood vessel density around developing follicles during the ovulatory stages, immunohistochemistry against CD31 was performed (marked with a white dashed line in the image insert, Figure 1j, k), and staining intensity was quantified (Figure 1l). The coverage of CD31 positive cells in the thecal layer is 4-fold higher during the estrus stage compared to the diestrus stage (Figure 1l, *p*<0.0001). The increased vascularity during the estrous cycle is restricted to the reproductive system, driven by vascular endothelial growth factor (VEGF) secretion^1^. Higher VEGF levels also lead to capillary leakiness and increased permeability, thus enabling nanoparticle extravasation through large endothelial gaps that are present during the proestrus and estrus stages^30^.

### Nanoparticle accumulation in the reproductive system is size-dependent

To test whether the ovaries have a nanoscale size cutoff, PEGylated gold nanoparticles (AuNPs) of different sizes: 20, 50, 100, and 200 nm were injected intravenously during the estrus stage^31^. Accumulation of AuNPs in the ovaries and the uterus was quantified 24 hours post-injection using elemental analysis (Figure 2a). After 24 hours, 2.1%±0.2 of the injected dose normalized to the tissue weight was quantified in the ovaries for 20 nm AuNPs and 1.9%±0.1 for 50 nm AuNPs were detected in the ovaries. Contrarily, only 0.9%±0.1 or 0.4%±0.02 of the 100 nm and 200 nm AuNP, respectively, reached the ovaries (Figure 2a). In a similar pattern, 20 and 50 nm AuNPs accumulated in higher doses at the uterus compared to 100 and 200 nm (Figure 2a). In summary, 100 nm AuNPs accumulated ~2-fold less in the ovaries (*p*<0.05) and ~3.5-fold less in the uterus (*p*<0.05) compared to smaller AuNPs sizes, while 200 nm AuNPs accumulated ~5-fold less in the ovaries (*p*<0.01) and ~12.5-fold less in the uterus (*p*<0.01) compared to smaller AuNPs sizes.

**Figure 2.**
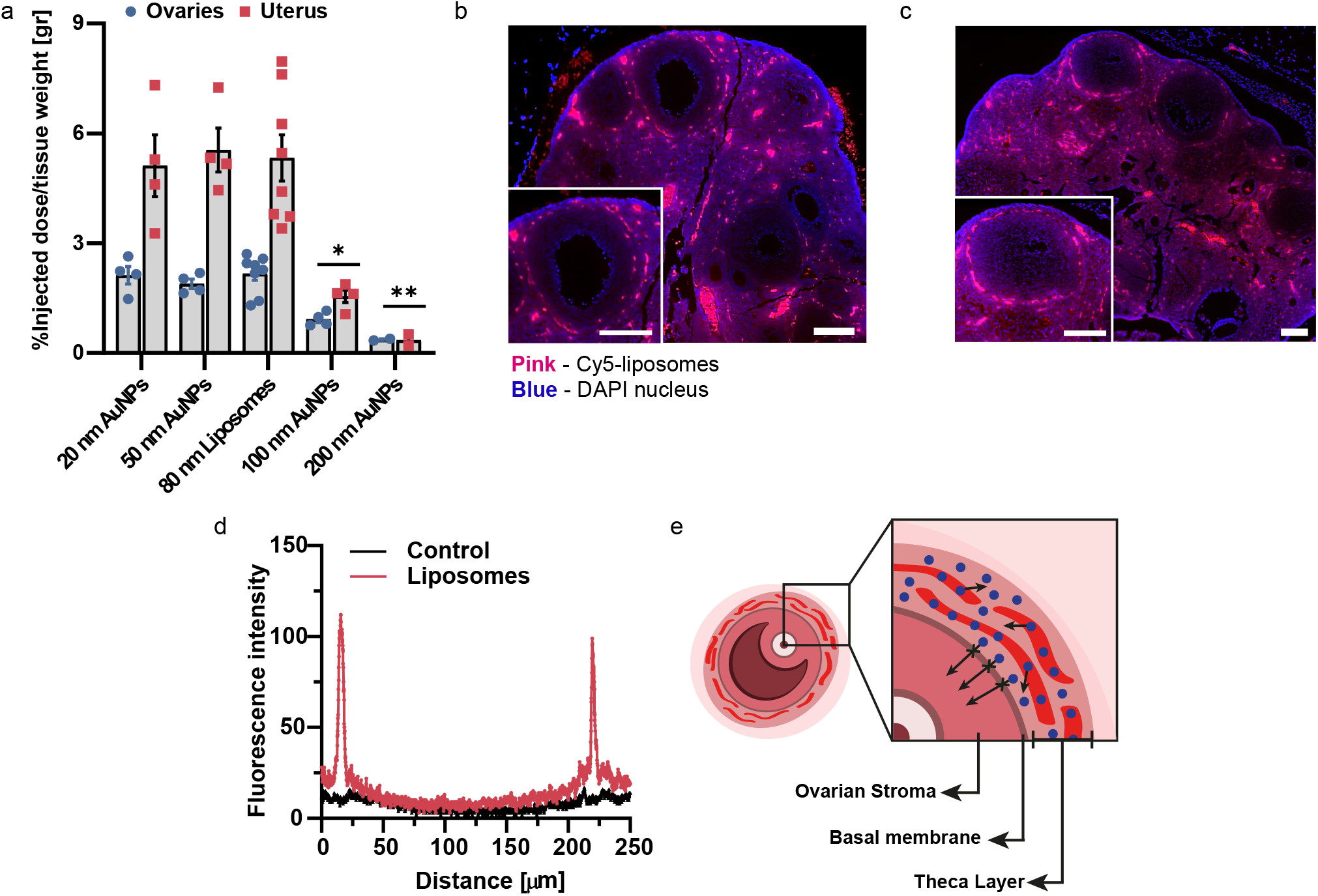
Nanoparticles are localized around the follicles with size-dependent accumulation in the reproductive system. Nanoparticles, 80-nm and smaller, show higher accumulation in the reproductive system, demonstrated by size dependent accumulation of gold nanoparticles in the ovaries (blue circles) and the uterus (red squares) 24 hour after i.v. injection during the estrus stage (n;4 for 20 nm, 50 nm and 100 nm; n;9 for 80 nm) (a). 80-nm liposomes do not cross the blood-follicle barrier as demonstrated by fluorescent histology images of cy5-labeled liposomes (pink) localization in the ovary (nuclei - blue) 24 hours after i.v. injection (b, c scale bar - 100 μm). Line profile of the fluorescent intensity signal across a single follicle shows that the liposomes surround the follicle, indicated by two peaks in the dye signal (d). Illustration of the blood-follicle barrier shows the liposomes (blue) do not cross the basal membrane of the follicle and are restricted to the thecal layer around the follicle (e). Results are shown as mean±SEM. Two-way ANOVA and Tukey’s t-test were used for statistical analysis of e. *p<0.05, **p<0.01.

### 80-nm liposomes are restricted outside the blood-follicle barrier

The blood-follicle barrier (BFB) forms during the early follicle stage and remains as a protective biological barrier until ovulation, at which point the barrier ruptures to release the oocyte^32^. Changes in the BFB structure throughout the follicular development allow strict regulation of the follicular fluid composition. The BFB protects the developing oocyte from toxic and foreign molecules while supplying it with necessary nutrients and growth factors^33^. 24 hours post i.v. injection, 80-nm liposomes were found surrounding the follicle, specifically at the outer thecal layer (Figure 2b, c). Fluorescence intensity measurements across a single follicle demonstrated that the liposomal signal is restricted outside the follicle, signified by two distinct peaks, with baseline signal recorded inside the follicle itself (Figure 2d), suggesting that 80 nm liposomes do not cross the BFB (Figure 2e).

### Doxorubicin-loaded liposomes cause delayed ovarian toxicity

We evaluated the effect of liposomal doxorubicin versus free doxorubicin on ovarian toxicity^13^. Healthy female mice received an i.v. injection of either free doxorubicin (free-DOX) or doxorubicin-loaded liposomes (DOX-lipo) during the estrus and diestrus stages (Figure 3a). After 24 or 48 hours, the ovaries were collected and RT-PCR analysis of pro- and anti-apoptotic genes was performed. Ovarian toxicity was recorded in all the experimental groups compared to the non-treated control (Figure 3b). The highest apoptosis level was measured 24 hours after free-DOX injection during estrus (R=2.7 ± 0.2, n=3, ratio between pro and anti-apoptotic gene expression normalized to healthy mice), and was significantly higher (*p*<0.05) compared to injection during the diestrus stage (R=1.5 ± 0.08, n=2). Furthermore, apoptosis levels 24 hours after injection at the estrus stage were ~1.5-fold higher for the free-DOX group compared to the DOX-lipo group (*p*<0.05). Immunohistochemistry analysis confirmed that the apoptotic effect took place inside the follicles (Figure 3c, d). Anti-active caspase3 staining was used to detect apoptotic follicles 4, 24, and 48 hours after injection of either free-DOX or DOX-lipo injected mice. At all time-points, the larger the follicle the greater the apoptosis levels were (Figure S1b, S1c). This agrees with the fact of increased blood supply to larger follicles^34^, and therefore increased drug exposure. After 24 hours, free-DOX treated mice had significant apoptosis in both small and large follicles (Figure 3c), compared to the DOX-lipo treated ovaries which showed no apoptotic signs in the smaller follicles (Figure 3d). The overall percentage of apoptotic follicles (Figure 3e) after 4 hours was slightly higher for the free-DOX treated ovaries (36% ± 5.6, n=5) compared to the DOX-lipo group (28% ± 3.3, n=3). After 24 hours, the percentage of overall apoptotic follicles increased significantly (*p*<0.0001) for the free-DOX treated group (64% ± 10, n;4) compared to the DOX-lipo group, which remained constant (32% ± 10, n=5). Surprisingly, after 48 hours, there was no significant increase in apoptosis in the free-DOX group (69% ± 9, n=5) while the percentage of apoptotic follicles spiked significantly (*p*<0.0001) in the ovaries of DOX-lipo treated mice (70% ± 5, n=5). Free-DOX was previously shown to cross the BFB and damage the developing oocytes^13,35^. These results suggest the delayed toxic effect of the DOX-lipo is due to initial liposomal accumulation in the ovaries, followed by drug release that induces the toxicity.

**Figure 3.**
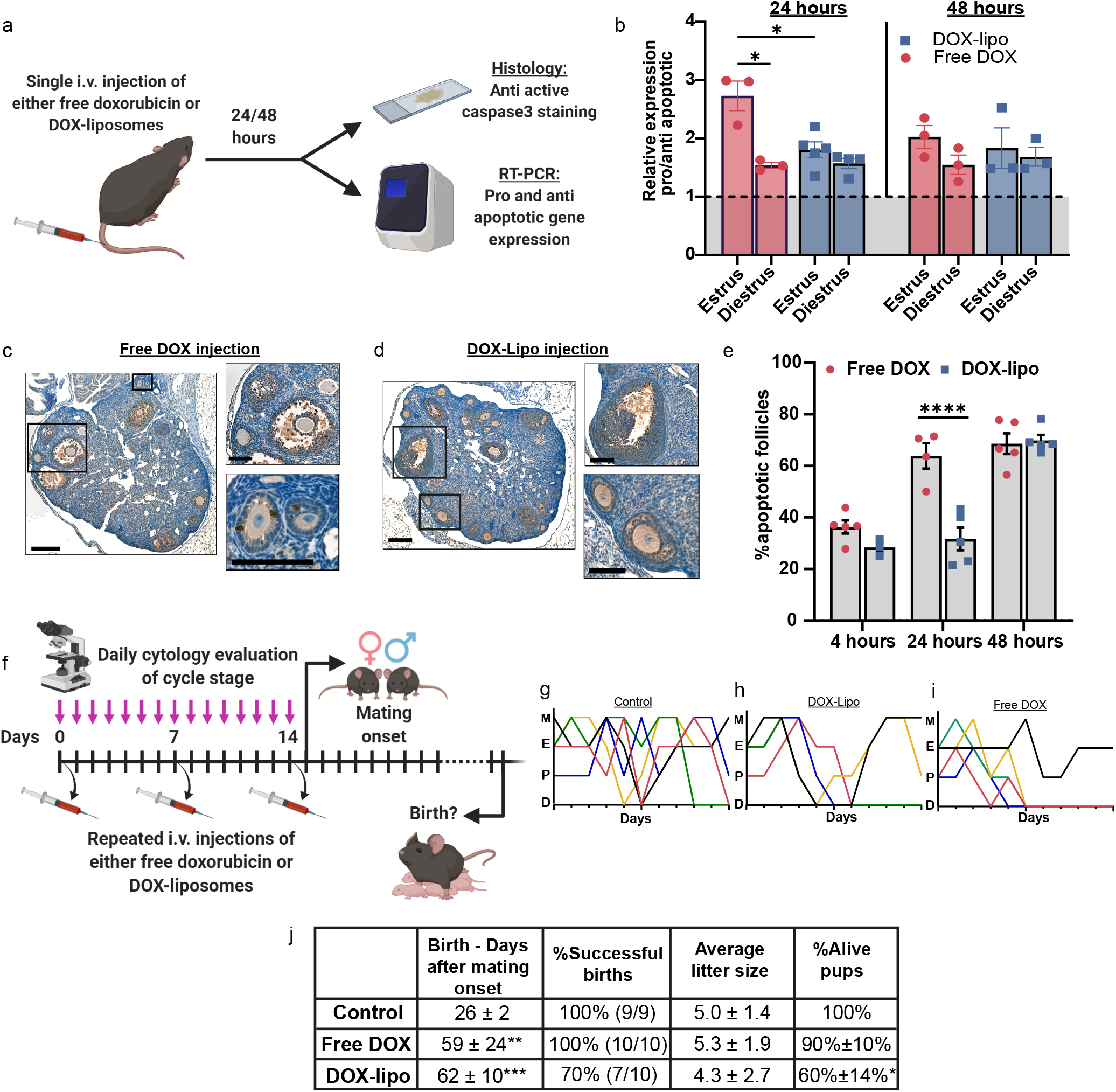
Doxorubicin-loaded liposomes cause delayed ovarian toxicity and impair fertility compared to the free drug. Healthy female mice received i.v. injection of either free DOX or DOX-lipo and the ovaries were taken to histology and RT-PCR analysis after 24 and 48 hours (a). RT-PCR of pro/anti apoptotic gene expression during estrus (24 hr, n;3 for free DOX n=5 for DOX-lipo. 48 hr, n;3 for both groups) and diestrus (24 hr, n=3 for free DOX n=4 for DOX-lipo. 48 hr, n=3 for both groups) are shown as the relative expression of the ratio between pro (BAX) and anti (bcl2) apoptotic genes (b). Immunohistochemistry of anti-active caspase3 24 hours after i.v. injection of either free DOX (c) or DOX-lipo (d) show apoptotic follicles (brown signal) with higher signal in the follicles in the free DOX group. Quantification of the total signal output from the follicles demonstrated significantly more follicles were damaged in the free DOX group (n=4) compared to DOX-lipo (n=5) 24 hours after i.v. injection, however apoptosis levels become comparable after 48 hours (n=5 for both groups) (e). Healthy female mice received repeated i.v. injections of either free DOX (n=10) or DOX-lipo (n=10) once a week for 3 treatments and their cycle stage was recorded daily to be compared to that of the control group (n=9) that was not injected (f). The estrous cycle in each group is demonstrated in graphs G-I where each line represents a single mouse. Mice in the DOX-lipo (h) and the free DOX (i) groups had a disrupted cycle compared to the control group (g). After the third injection, females were housed with males and the day of birth, litter size and pups’ viability were recorded (j). The time until pregnancy was significantly longer for both the free DOX and the DOX-lipo group, however the pups’ viability was lower for the DOX-lipo group. Results are shown as mean±SEM. Three-way ANOVA and Tukey’s t-test (b), two-way ANOVA and Tukey’s t-test (e and %alive pups in j) and Dunnett’s T3 t-test (days until birth in j) were used for statistical analysis. *p<0.05, **p<0.01, ***p<0.001, ****p<0.0001.

### Treatment with doxorubicin-loaded liposomes impairs fertility compared to the free drug

We further explored the effect of nanoparticulate doxorubicin on fertility. Healthy female mice were divided into three groups: free-DOX treated, DOX-lipo treated, and non-treated control. The two treatment groups received three rounds of weekly i.v. injections and their cycle stage was monitored daily (Figure 3f). After three treatments, the females were housed with male mice for mating. The day of birth was monitored in addition to the litter size, pups’ weights and viability after birth. The toxic effect of both DOX and DOX-lipo was apparent during the treatments as the estrous cycle was disrupted in the DOX-lipo treated group (Figure 3h) and the free-DOX treated group (Figure 3i), compared to the control group where the estrous cycle remained normal. All mice (100%) in the control group had successful births 26 ± 2 (n=9) days after mating onset (Figure 3j). In contrast, time until first litter was 59±24 days (n=10) for the free-DOX treated group and 62±10 days (n=10) in DOX-lipo group. Furthermore, while all mice in the free-DOX group were pregnant and gave birth (100% births), only 70% of the mice in the DOX-lipo group conceived. This implies that the mice in the DOX group regained their ability to ovulate. The average litter size was 5±1.4 pups, 5.3±1.9, and 4.3±2.7 for the control, free-DOX, and DOX-lipo group, respectively. The viability of the pups was 100% and 90%±10% for the control and the free-DOX groups respectively while it declined to merely 60%±14% for the DOX-lipo group (*p*<0.05). These results suggest that the treatment with DOX-lipo increases ovarian toxicity compared to free-DOX. This can be explained by the long retention time of the liposomes in the ovaries compared to free molecule drugs that are cleared rapidly^36^.

### The estrous cycle affects tumor biodistribution and nanomedicine therapy

Nanomedicine treatments are used as first-line therapies against several types of cancer^37,38^. We tested the effect of the ovulatory cycle on nanoparticle accumulation in tumors and the reproductive system. For this, we measured the biodistribution of 80-nm liposomes during estrus and diestrus stages in mice bearing orthotopic triple-negative breast cancer (4T1) tumors (Figure 4a) or epithelial ovarian cancer (Figure 4b). Gd-lipo, 80 nm, were i.v. injected to tumor-bearing mice and their accumulation was quantified 24 hours after injection. In the breast cancer model, 2.6-fold more liposomes accumulated in the reproductive system compared to the tumor during the estrus stage (*p*<0.05). Interestingly, 4.1-fold more liposomes accumulated at the tumor compared to the reproductive system during the diestrus stage (*p*<0.01), indicating an opposite trend. These results suggest that during the estrus stage, the accumulation of nanoparticles is shifted towards the reproductive system rather than the tumor (Figure 4a).

**Figure 4.**
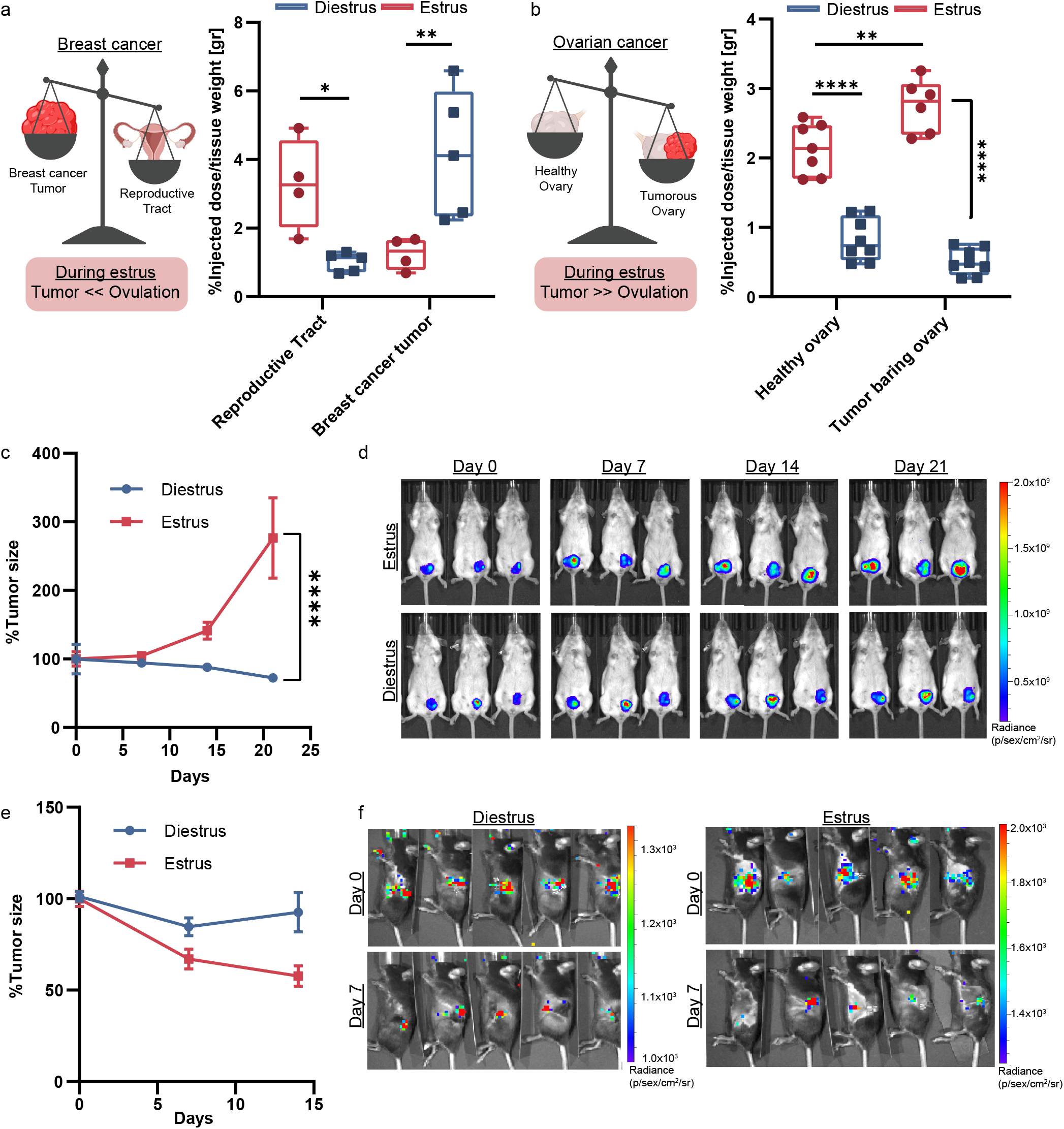
Biodistribution of liposomes during cancer and cancer treatment efficacy are affected by the female mouse cycle. During breast cancer tumor treatment, Gd-liposomes accumulate in the reproductive system (n;4) instead of at the site of the tumor (n;4) during the estrus stage, while during the diestrus stage they shift towards the tumor (n;5) and away from the reproductive system (n;5) (a). During ovarian cancer, there is increased accumulation of Gd-liposomes both in the tumor-bearing ovary (E. n;6, D. n;8) and in the healthy ovary (E. n;6, D. n;7) during estrus, compared to during diestrus stages (b). Treatment efficacy of DOX-lipo was evaluated in 4T1 mcherry breast cancer model during estrus (n;7) and diestrus (n;6) (c). IVIS images of 4T1 mcherry breast cancer tumor (top - treatment during estrus, bottom - treatment during diestrus) show 3 representative mice from each group (d). Treatment efficacy of DOX-lipo was evaluated in an ovarian cancer model expressing luciferase during estrus (n;5 (days 0 and 7), n;2 (day 14)) and diestrus (n;5 (days 0 and 7), n;3 (day 14)) (e). IVIS images of luminescent ovarian cancer tumor, left panel - treatment during diestrus, right panel - treatment during estrus (f). Results are shown as mean±SEM. Two-way ANOVA and Tukey’s t-test were used for statistical analysis of a-c, e. **p<0.01, ***p<0.001, ****p<0.0001.

In the case of ovarian cancer, 2.7%±0.15 (n;6) of the injected dose accumulated at the tumor-bearing ovary while 2.1%±0.14 (n;7) reached the neighboring healthy ovary at the time of estrus. This 1.3-fold difference (*p*<0.01) implies that cancer vascularization in the tumor-bearing ovary overcomes ovulation angiogenesis in the healthy ovary during the estrus stage.

We then tested the efficacy of DOX-lipo treatment on these cancer models during estrus and diestrus stages. Triple-negative breast cancer tumor-bearing mice were treated weekly with an intravenous administration of DOX-lipo, either during the estrus stage or diestrus stage, for three treatment cycles (Figure 4c, d). After 21 days, the average tumor size in the estrus group increased by 276%±58 compared to day 0, while in the diestrus-treated group the tumor reduced to 72%±6 of the initial size (*p*<0.0001, Figure 4c). This result correlates with the biodistribution profile during the estrus stage, in which the nanomedicine preferentially accumulated in the reproductive system rather than at the tumor, resulting in decreased therapeutic efficacy.

When the reproductive system is the target of the nanomedicine treatment, as in ovarian cancer, the outcome is different. Treatment of ovarian cancer with DOX-lipo was favorable during the estrus stage compared to the diestrus stage (Figure 4e, f). After 14 days, the average tumor size of the diestrus group was ~1.6-fold higher than the average tumor size of the estrus group, as more liposomes reach the tumor-bearing ovary during ovulation thus increasing efficacy. This finding can be leveraged as a treatment strategy to increase nanomedicine accumulation in ovarian tumors for women that are still of reproductive age, while also accounting for possible effects on fertility.

## Conclusions

Our findings show that the menstrual cycle affects the biodistribution of therapeutic nanoparticles towards the female reproductive system. Proper sex considerations in preclinical studies and clinical trials are encouraged^39^, however in the field of nanotechnology these considerations are still nascent. Nanoparticle accumulation during the estrous cycle should be considered in future studies, as it may lead to increased variability in biodistribution and drug efficacy. Our findings on possible ovarian toxicity caused by DOX-lipo compared to free-DOX should be taken into consideration when devising a treatment plan for patients in their reproductive age. These findings may also be leveraged for designing targeted treatments of the ovarian follicles for controlled drug release in this essential tissue of life.

## Acknowledgments

This work was supported by ERC-STG-2015-680242.

Maria Poley wishes to thank the Israeli Ministry of Science and Technology for the Shulamit Aloni Doctoral Fellowship. Omer Adir wishes to thank the Miriam and Aaron Gutwirth Memorial Fellowship. Maya Kaduri wishes to thank TEVA Pharmaceuticals - NFBI - The National Forum for BioInnovators for a doctoral grant and the Technion Integrated Cancer Center (TICC) Rubinstein scholarship. The authors also acknowledge the support of the Technion Integrated Cancer Center (TICC), the Russell Berrie Nanotechnology Institute, the Lorry I. Lokey Interdisciplinary Center for Life Sciences & Engineering, The Israel Ministry of Economy for a Kamin Grant (52752, 69230); the Israel Ministry of Science Technology and Space – Office of the Chief Scientist (3-11878); Israel Innovation Authority for Nofar Grant (67967), the Israel Science Foundation (1778/13, 1421/17); the Israel Ministry of Science & Technology (3-16963, and Covid-19 grant); the Israel Cancer Association (2015-0116); the German-Israeli Foundation for Scientific Research and Development for a GIF Young grant (I-2328-1139.10/2012); the European Union FP-7 IRG Program for a Career Integration Grant (908049); the Phospholipid Research Center Grant; the Louis family Cancer Research Fund, a Mallat Family Foundation Grant; The Unger Family Fund; a Carrie Rosenblatt Foundation for Cancer Research, A. Schroeder acknowledges Alon and Taub Fellowships. Images in this paper were created with BioRender.com and Adobe Illustrator.

## Materials and Methods

### Estrous cycle and cytology

Healthy 8-10 week old C57BL/6 female mice were housed 5 per cage in standard laboratory conditions and were exposed to 12 hours of light and 12 hours of dark. All animal experiments were approved by, and in compliance with, the institutional and ethical committee at the Technion. The animals’ well-being was monitored regularly. Ovulation and cycle synchronization was achieved by mixing solid bedding from the cages of male mice with the bedding of the female mice cage for 2 days^40^. The estrous cycle stage of each female mouse was determined using the cytology method which was performed daily at the same time (7AM-9AM). Vaginal cytology is a common method for cycle stage evaluation by simply observing the different cell populations in stained vaginal smears taken from the vaginal opening of the mouse^41^. To collect a vaginal smear, 10 μl of sterile water were carefully pipetted on the vaginal opening and then placed on a glass slide to air dry. The dry smears were stained using Jorvet Dip Quick staining kit (Jorgensen Laboratories) and examined under a light microscope. The cycle stage was determined by observing the cell population (Figure S1A). Before experiment initiation, two consecutive estrous cycles were confirmed for each mouse.

### Gadolinium loaded liposomes preparation

Gd liposomes were prepared as described before^42^. Briefly, a lipid mixture of hydrogenated soybean phosphatidylcholine (HSPC; Avanti Polar Lipids, Alabaster, AL, USA), cholesterol (Sigma-Aldrich, Rehovot, Israel) and 1,2-distearoyl-sn-glycero-3-phosphoethanolamine-N-methoxy-polyethylene glycol 2000 (DSPE-PEG2000; Avanti Polar Lipids, Alabaster, AL, USA), in molar percentages of 56:39:5 was dissolved in pure ethanol at 70°C. The lipid mixture was injected into a Dulbecco’s Phosphate Buffer Saline (PBS; Sigma-Aldrich, St. Louis, USA) solution containing 167 mg/ml of Gd-DTPA Diethylenetriaminepentaacetic acid gadolinium(III) dihydrogen salt hydrate (Gd; Sigma-Aldrich, Rehovot, Israel) to obtain a final lipid concentration of 50 mM. The liposomes were downsized to 80 nm using a Lipex extruder (Northern Lipids, Vancouver, Canada) at 65°C through 400, 200, 100, 80 and 50 nm Nuclepore polycarbonate membranes (Whatman, Newton, MA, USA). Free Gd-DTPA was removed using dialysis in a 12-14 kD membrane (Spectrum Laboratories, Inc., USA) against PBS (1:1000 volume ratio) at 4°C and exchanged three times. Average liposome size was measured using Zetasizer Nano ZSP (Malvern Instruments, UK) in disposable polystyrene cuvette after liposomes were diluted 1:100 in PBS and Cryo-TEM was performed as described previously^42^.

### Cy5-labeled liposomes preparation

Cy5-labeled liposomes were prepared in the same method as Gd-liposomes only without Gd added to the PBS solution. 1% molar percentage of DSPE-Cy5 was added to the lipid mixture before injection to PBS solution.

### Doxorubicin loaded liposomes preparation

Doxorubicin (DOX, TEVA Israel) was actively loaded into a 80 nm liposome using the ammonium sulfate gradient method^43^. Lipid mixture of HSPC, cholesterol and DSPE-PEG2000 in molar percentages of 56:39:5 respectively was dissolved in pure ethanol at 70°C. The dissolved lipids were injected into 120 mM ammonium sulfate solution to reach a final concentration of 50 mM total lipids. The liposomes were downsized to 80 nm using an extruder at 70°C. Dialysis was performed in 12-14 kD dialysis membrane against 10% w/w sucrose and 10 mM histidine at pH 6.5 (1:1000 volume ratio) and exchanged three times. For active loading, DOX was dissolved in 10% w/w sucrose and added to the ammonium sulfate liposomes to reach a final concentration of 2 mg/ml. The mixture was placed in 70°C at 600 rpm for 1 hour. The dox loaded liposomes were dialyzed in 12-14 kD dialysis membrane against 10% w/w sucrose and 10 mM histidine pH 6.5 (1:1000 volume ratio) for 24 hours.

### Gd-liposomes biodistribution

After determining the estrous cycle, 200 μl of Gd-liposomes were injected i.v. into the tail vein of healthy female mice. 24 hours after injection, the mice were sacrificed and the ovaries, uterus, kidneys, spleen, liver, heart, and lungs were collected and weighed. The organs were disintegrated to ash at 550°C for 5 hours. The ashes were dissolved in 5 ml of 1% nitric acid and the mixture was filtered through 0.45 μm syringe filters and taken for ICP-OES analysis.

### Gold nanoparticles biodistribution

20, 50 and 100 nm PEGylated gold nanoparticles were purchased from NanoCompasix (San Diego, CA, USA), and 200 nm PEGylated gold nanoparticles were purchased from NanoPartz (Loveland, CO, USA). After determining the estrous cycle, 100 μl of 1 mg/ml gold nanoparticles were injected i.v. into the tail vein of healthy female mice. 24 hours after injection the mice were sacrificed and the ovaries and uterus were collected and weighed. The organs were dissolved overnight in Aqua-regia solution and then heated at 60°C for 1 hour. The dissolved organs were resuspended in double distilled water to reach acid concentration of 1%. The mixture was filtered through 0.45 μm syringe filters and taken for ICP-OES analysis.

### Elemental analysis of Gd and Au

Gadolinium (Gd) and Gold (Au) samples were analyzed using inductive coupled plasma – optical emission spectroscopy 5110 ICP-OES (Agilent, CA, USA). Calibration curves for each element were obtained from a calibration standard (Sigma-Aldrich, Rehovot, Israel) diluted in 1% nitric acid. Gd emission was measured at 335.048 nm and 342.246 nm. Au emission was measured at 242.794 nm and 267.594 nm. The concentration at each wavelength was calculated by the ICP-OES software according to the obtained calibration curve. The measurements from both wavelengths were averaged for each element. The obtained concentration was divided by the injected dose concentration to obtain the percentage out of the injected dose and then divided by the organ’s weight for normalization.

### *Ex-vivo* IVIS imaging

200 μl of Cy5 labeled liposomes were injected i.v. into the tail vein of healthy female mice after determining the estrous cycle stage. 24 hours after injection the mice were sacrificed and the reproductive system was imaged *ex-vivo* using the IVIS SpectrumCT Pre-Clinical *In-Vivo* Imaging System (PerkinElmer, MA, USA) at an excitation of 570 nm and emission of 620 nm, binning 8, f-stop 2 and 3 seconds exposure. A control (not-injected) mouse was used for analysis. All images were analyzed using the LivingImage software.

### Fluorescent Histology analysis

200 μl of Cy5 labeled liposomes were injected i.v. into the tail vein of healthy female mice after the estrous cycle stage was determined. 24 hours after injection the mice were sacrificed and the reproductive system was fixated using Formalin solution neutral buffered 10% histological tissue fixative (Sigma-Aldrich, Rehovot, Israel) at 4°C for at least 24 hour before embedding in paraffin and sectioned. Slides were deparaffinized in a xylene ethanol gradient as follows: soaked in xylene for 3 min, xylene/ethanol (1:1 vol ratio) for 3 min, absolute ethanol for 3 min, 95% ethanol for 3 min, 70% ethanol for 3 min, and 50% ethanol for 3 min, and finally placed in tap water. Nuclei blue fluorescent staining was done using Invitrogen™ Molecular Probes™ NucBlue Fixed Cell ReadyProbes™ Reagent (Fisher Scientific, Waltham, MA, USA) incubated for 5 min followed by rinsing with tap water. The slides were imaged with Leica DMI8 inverted fluorescent microscope (Leica Microsystems GmbH, Wetzlar, Germany) using x40 magnification with exposure times of 400 ms for the DAPI channel and 900 ms for the Cy5 channel.

### Immunohistochemistry analysis

Slides were deparaffinized in a xylene ethanol gradient as follows: soaked in xylene for 3 min, xylene/ethanol (1:1 vol ratio) for 3 min, absolute ethanol for 3 min, 95% ethanol for 3 min, 70% ethanol for 3 min, and 50% ethanol for 3 min, and finally placed in tap water. Antigen retrieval was done in 10 mM tri-sodium citrate solution at pH 6 tittered with HCl. 2.5% ready-to-use normal goat serum (vector laboratories) was used for blocking. Incubation with primary antibody was at 4°C overnight with either anti-CD31 diluted 1:100 (ab28364, Abcam, Cambridge, MA, USA) or anti-active caspase3 diluted 1:100 (ab2302, Abcam, Cambridge, MA, USA). For blocking of endogenous peroxidase activity, the slides were incubated for 30 minutes in 0.3% hydrogen peroxide solution and then washed in tap water before incubation with ready-to-use secondary goat anti-rabbit antibody conjugated to HRP (MP7451 kit, vector laboratories) for 40 minutes at room temperature. For color development, slides were incubated with DAB solution (SK4105 kit, vector laboratories) for 3 minutes, washed with tap water, and counter stained with hematoxylin. Slides were scanned using 3DHistech Panoramic 250 Flash III automated slide scanner (3DHistech, Budapest, Hungary) at x40 magnification.

### Apoptosis experiment

Healthy female mice were divided into 3 groups: control, Dox-liposomes and free DOX. After determining the estrous cycle stage, mice were injected with 100 μl of 5 mg/kg of either Dox-liposomes or free DOX. The mice were sacrificed either after 24 or 48 hours and the reproductive system was taken for either mRNA extraction for RT-PCR or fixation in 10% Formalin solution neutral buffered histological tissue fixative for immunohistochemistry analysis.

### RT-PCR analysis

mRNA was extracted from ovaries using NucleoSpin RNA kit (#740955.50, Macherey-Nagel, Germany) according to the manufacturer’s protocol. mRNA was quantified by measuring absorbance at 260 nm with a NanoQuant plate in an Infinite 200PRO plate reader (TECAN, Mannedorf, Switzerland). 400 ng of mRNA were used for cDNA synthesis in a 20 μl reaction volume using high-quality cDNA synthesis kit (PCR Biosystems, Wayne, PA, USA). Real-time PCR was performed on qRTPCR CFX Bio-Rad machine (Bio-Rad laboratories, Hercules, CA, USA) using qPCRBIO Fast qPCR SyGreen Blue Mix Hi-ROX (PCR Biosystems, Wayne, PA, USA). GAPDH gene was used as a house keeping gene for normalization. Primers sequences:

Bcl2-Forward-ATGCCTTTGTGGAACTATATGGC Reverse-GGTATGCACCCAGAGTGATGC, BAX-Forward-TGAAGACAGGGGCCTTTTTG Reverse-AATTCGCCGGAGACACTCG, GAPDH-Forward-TGCACCACCAACTGTTAG Reverse-GGATGCAGGGATGATGTTC.

### Mating experiment

Healthy 8-10 weeks old C57BL/6 female mice were divided into 3 groups: control, free DOX and Dox-liposomes. The estrous cycle of all the mice was monitored daily. 5 mg/kg of either free DOX of Dox-liposomes were injected i.v. into the mice tail vein every 7 days for total of three injections per group (Day 0, 7 and 14). After the third injection, 2 females were placed with a healthy C57BL/6 male and were allowed to mate. Pregnancy was monitored and the day of birth, number, viability and weight of pups were documented.

### Cell culture

MOSE cells were kindly provided by Novocure Ltd. Triple negative breast cancer cells 4T1 mCherry cells were purchased from ATCC. Cells were mycoplasma free. MOSE-luc cells were cultured in Dulbecco’s Modified Eagle Medium: Nutrient Mixture F-12 (DMEM-F12, Biological Industries, Israel) supplemented with 10% fetal bovine serum (FBS), 100 IU ml-1 Penicillin, 100 μg ml-1 Streptomycin and 2 mM L-glutamine (Biological Industries) and grown at 37°C; 5% CO2. 4T1 mCherry cells were cultured in Roswell Park Memorial Institute medium (RPMI; Sigma-Aldrich, Rehovot, Israel) supplemented with 10% FBS, 100 IU ml-1 Penicillin, 100 μg ml-1 Streptomycin and 2 mM L-glutamine and grown at 37°C; 5% CO2. To detach the cells, they were first incubated in fresh medium for 1 hour, washed with PBS and incubated with trypsin for 5 minutes. The cells were collected by centrifugation at 200Xg for 5 minutes. The cell pellet was suspended in fresh medium and cell concentration was measured using a cell counter.

### Breast cancer model

50 μl of 250,000 4T1 mCherry cells were suspended in PBS and injected subcutaneously directly into the mammary fat pad of healthy 8-10 weeks old BALB/c female mice^44^. Tumor development was monitored using caliper measurements and IVIS *in-vivo* fluorescent imaging. The mice were treated either during estrus or diestrus stages with 6 mg/kg of Dox-liposomes once a week for 21 days.

### Orthotopic ovarian cancer model

Healthy 8-10 week old C57BL/6 female mice were anesthetized using a mixture of 90 mg/kg ketamine and 10 mg/kg of xylazine and received 0.3 mg/kg of buprenorphine for pain relief. Small incisions in the skin and peritoneum were made in the back of the animal above the ovarian fat pad. The ovary was pulled out and held in place with a bulldog clip by the ovarian fat pad. Using a 30-gauge Hamilton syringe 10 μl of 100,000 MOSE (expressing luciferase) cells suspended in equal parts of PBS and Matrigel were injected directly into the ovarian bursa. The ovary was placed gently back inside and the peritoneum was sutured using 7-0 absorbable stitches. The skin was closed using surgical clips. The animals were placed in a 37°C incubator until fully awake. Tumor development was monitored using D-luciferin injection (150 mg/kg) followed by IVIS *in-vivo* luminescence imaging. The mice were treated either during estrus or diestrus stages with 6 mg/kg of Dox-liposomes once a week for 14 days.

### *In-vivo* IVIS imaging

Whole animal imaging was performed in the IVIS Spectrum CT Pre-Clinical *In-Vivo* Imaging System (PerkinElmer, MA, USA). For all imaging, the animals were placed under isoflurane anesthesia. 4T1 mCherry tumors were imaged using fluorescent mCherry filter with the following settings: ex. 570 nm em. 620 nm, binning 4, f-stop 2 and exposure time 3 seconds. Ovarian luciferase expressing tumors were imaged using the luminescence setting with the following settings: open emission filter, binning 16, f-stop 1 and exposure time of 180 seconds, 12 minutes after i.p injection of 150 mg/kg of D-luciferin. All images were analyzed using the LivingImage software.

## Statistical analysis

All statistical analysis: student’s t-test, two-way ANOVA and three-way ANOVA were performed using Prism GraphPad software.

