## Supplemental figure for "Chemotherapeutic Nanoparticles Accumulate in the Female Reproductive System during Ovulation Affecting Fertility and Anticancer Activity"

Supplementary figure

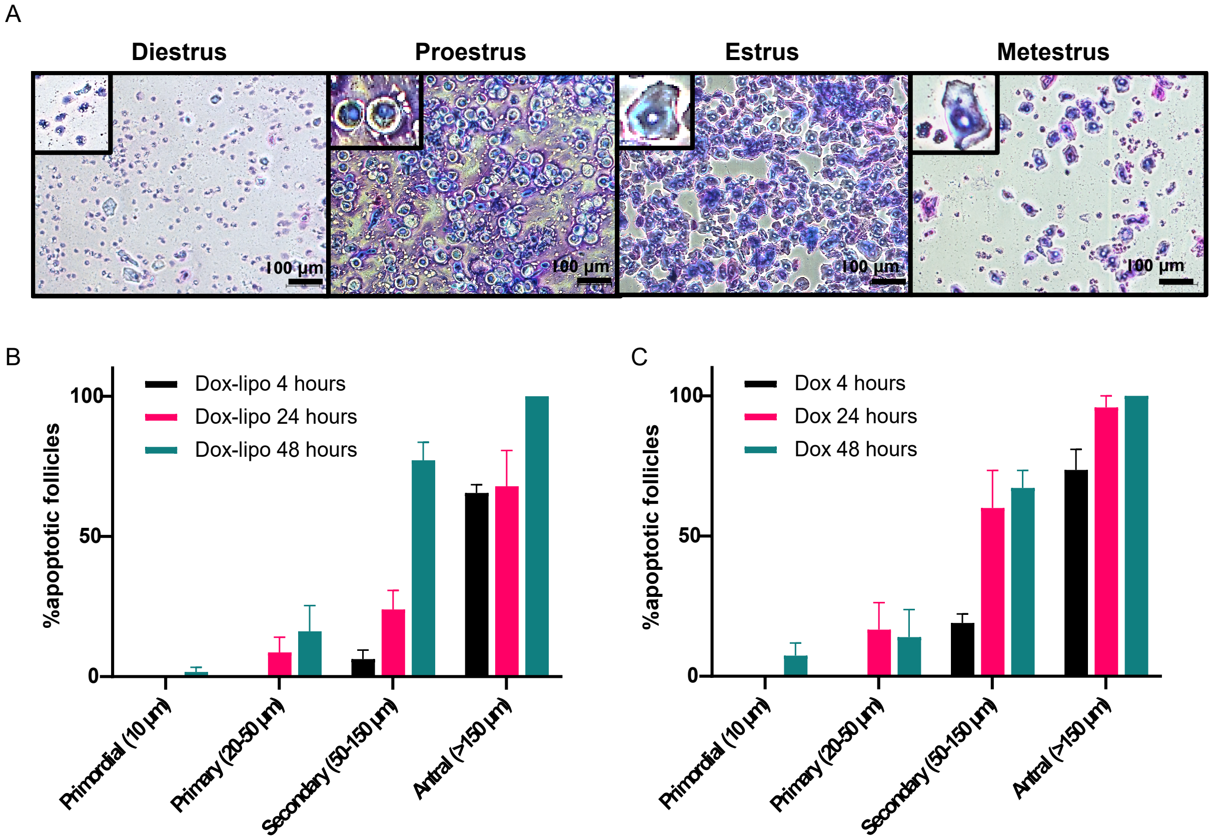


**Figure S1.** Cytology smears at different cycle stages of a healthy female C57BL/6 mouse. Enlarged imaged of each representative cell type is shown in the top left corner. Scale bar 100 µm (A). Percentage of apoptotic follicles categorized by their size 4, 24, and 48 hours after Dox-lipo i.v. administration (B).Percentage of apoptotic follicles categorized by their size 4, 24, and 48 hours after free-DOX i.v. administration (C).
